# Loss of charge at solvent exposed Lys residues does not induce the aggregation of superoxide dismutase 1

**DOI:** 10.1101/301309

**Authors:** Keith Crosby, Anthony M. Crown, Brittany L. Roberts, Hilda Brown, Jacob I. Ayers, David R. Borchelt

**Author notes:** These authors contributed equally to the content of this paper and are presented in alphabetical order (Keith Crosby, Anthony M. Crown). To whom correspondence should be addressed pertaining to structural analyses: David R. Borchelt, Department of Neuroscience, Box 100159, 1275 Center Drive, University of Florida, Gainesville, FL 32610.

## Abstract

Mutations in superoxide dismutase 1 (SOD1) associated with familial amyotrophic lateral sclerosis (fALS) induce the protein to misfold and aggregate. To date, missense mutations at more than 80 different amino acid positions have been associated with disease. How these mutations perturb native structure to heighten the propensity to misfold and aggregate is unclear. One potential mechanism that has been suggested is that when mutations occur at positions occupied by charged amino acids, then repulsive forces that would inhibit aberrant protein:protein interactions would be reduced. Mutations at twenty-one charged residues in SOD1 have been associated with fALS. Here, we examined whether loss of positively charged surface Lys residues would induce the misfolding and aggregation of SOD1. We randomly mutated four different Lys residues (K30, K36, K75, K91) in SOD1 and expressed these variants as fusion proteins with yellow fluorescent protein (YFP). We also assessed whether these mutations induced binding to a conformation-restricted SOD1 antibody, designated C4F6, which recognizes non-natively folded protein. Our findings indicate that SOD1 generally tolerates mutations at surface exposed lysine residues, and that loss of positive charge is insufficient to induce aggregation. Our findings may explain why mutations at these Lys residues have not been identified in ALS patients.

## INTRODUCTION

Amyotrophic Lateral Sclerosis (ALS) is a fatal neurodegenerative disease primarily characterized by loss of upper and lower motor neurons. Although most forms of ALS are of unknown etiology (sporadic ALS), a subset of cases demonstrate dominant patterns of inheritance in specific proteins (familial ALS or fALS). Of these inherited genetic mutations, approximately 20% are found in Cu-Zn superoxide dismutase (SOD1) [1], the ubiquitous antioxidant protein responsible for detoxifying oxygen radicals in the cytoplasm [2,3]. SOD1 is a homodimer composed of 153-amino acid subunits in which each subunit contains eight β-strands, a catalytic copper ion, a structurally important zinc ion, an electrostatic loop element that forms a portion of the active site funnel, and an intramolecular disulfide bond between cysteine 57 and cysteine 146 [4–6]. Over 160 mutations in SOD1 have been associated with ALS {http://alsod.iop.kcl.ac.uk/default.aspx}. Disease onset for SOD1-fALS patients is 45-47 years [7], whereas the average age of onset in sALS cases tends to be later (55-60 years of age) [8].

The vast majority of SOD1 mutations associated with ALS are missense point mutations. The effects of fALS mutations on the normal enzyme activity and protein turnover vary greatly [9–13]. While some mutants are rapidly degraded or inactive, others retain high levels of activity and relatively long half-lives [9–18]. Mutant SOD1 of fALS is generally viewed as being more prone to misfold and aggregate [14,19–23]. SOD1 immuno-reactive inclusions in surviving spinal motor neurons is a common, but not uniformly found, pathologic feature of SOD1-linked fALS [24–39]. Notably, the SOD1 inclusions found in patients appear to lack the features of amyloid (Thioflavin and Congo Red negative) [24,40]. Misfolded SOD1 has also been described as a pathologic feature of sporadic ALS using antibodies that are preferentially reactive to non-natively folded SOD1 [41–43]. However, other studies have disputed these findings [44–46]. Thus, although the role of wild-type SOD1 in sporadic ALS requires further study, there is substantial evidence of misfolded and aggregated SOD1 in fALS patients with SOD1 mutations (reviewed in [47]).

In cell culture and mouse models of WT and mutant SOD1 expression, there is a clear distinction between the aggregation propensities of WT and mutant protein. Although mice expressing high levels of human WT SOD1 develop clinical signs of ALS, including paresis, the age at which WT mice reach endstage is approximately 2 times longer than mice that express comparable levels of G93A fALS SOD1 (367±56 days vs 155±9 days) [48]. At end-stage, the levels of mis-folded, aggregated SOD1 detected by filter-trap assay [49], were two-fold higher in paralyzed G93A mice than in paralyzed WT overexpressing mice [48]. Mice expressing WT-SOD1 fused to yellow fluorescent protein (SOD1:YFP) age normally and show little or no evidence of WT-SOD1:YFP aggregation, whereas equivalently expressed fALS mutant G85R-SOD1:YFP produces clinical signs of ALS with evidence of mutant protein oligomerization, aggregation, and inclusion formation [50]. In cell culture studies of SOD1 aggregation, we have consistently observed that WT SOD1 is >10-fold less prone to form detergent-insoluble aggregates than fALS mutant SOD1 (>40 mutants tested) [7,51,52]. These studies use a paradigm of transient over-expression in which aggregation of mutant SOD1 occurs over a period of 24 to 48 hours. The aggregates generated in these cell models are similar to what is generated in transgenic mouse models expressing mutant SOD in that in both cases the protein that acquires detergent-insolubility lacks a normal intramolecular disulfide bond [53].

Similar transient expression paradigms have been used to visualize the aggregation of mutant SOD1 in cultured cells, using a strategy in which SOD1 is fused to a fluorescent reporter protein [54–60]. In cells expressing mutant SOD1 fused to the fluorophore, fluorescent inclusions were observed, whereas WT SOD1 fused to fluorophore was diffusely distributed throughout the cytosol. We have demonstrated that WT and mutant SOD1 fusions proteins with YFP show differing propensities for inclusion formation, differences in detergent solubility, and differences in diffusibility in disrupted cells [61]. SOD1:YFP fusions with fALS mutations A4V, G37R, G85R, D101N, C111Y, S134N all readily form fluorescent inclusions when over-expressed [61,62]. Untagged versions of SOD1 with these same mutations form detergent insoluble, sedimentable, aggregates when transiently over-expressed [19,52]. Moreover, in studies that have experimentally combined familial and experimental mutations in SOD1 to examine the role of disulfide cross-linking in aggregation, we have observed complete agreement between the aggregation propensities of untagged and YFP tagged SOD1 variants [62]. Collectively, this body of work demonstrates that fALS mutations in SOD1 share a common feature of promoting aggregation of the protein, and that visualizing aggregation by expressing YFP fusion proteins is a useful approach.

The objective of the current study was to investigate whether loss of positively charged residues in SOD1 induce the aggregation of SOD1. It has been proposed that mutations that reduce overall charge may enhance the propensity for proteins to aggregate by reducing the repulsive forces that could occur when charged residues try to align and stack in aberrant aggregates, or when two monomers of misfolded protein interact in prelude to the formation of stronger aberrant interactions [63–65]. Our primary assay for aggregation in this effort was visualization of inclusions in cells over-expressing SOD1:YFP fusion proteins. In prior studies, we have used both HEK293 and Chinese Hamster Ovary (CHO) cells to visualize the aggregation of mutant SOD1 fused to YFP, finding similar results [61,62]. Here we have used the CHO model largely because these cells tend to be flat with a large cytosol that allows for a clearer assessment of aggregation. An important aspect of this approach is that the SOD1:YFP proteins are highly over-expressed. In this setting any modulation of aggregation by chaperone function or variabilities in protein stability are minimized, revealing inherent aggregation propensities of the protein [19,66]. In these over-expression model systems, both WT and mutant SOD1 are largely deficient in Cu ions and are less able to form the normal intramolecular disulfide bond associated with full maturation [66]. Thus, the model is essentially assessing the inherent propensity of immature mutant SOD1 to achieve native conformation. Our experimental data with this model indicates that loss of charge mutations in highly conserved Lys residues is not sufficient to induce SOD1 to spontaneously aggregate.

## RESULTS

In examining the frequency of mutations in positively charged residues of SOD1, we noted that only one of 11 Lys and one of four Arg residues are mutation sites in ALS {http://alsod.iop.kcl.ac.uk/default.aspx} (Fig. 1). Five of 7 His residues are sites of mutation, but these amino acids play critical roles in the binding of metal cofactors that contribute to the stability of native SOD1 structure [67]. By contrast, nine of 11 Asp and five of nine Glu negatively charged residues have been identified as sites of mutation in ALS {http://alsod.iop.kcl.ac.uk/default.aspx} (Fig. 1). To determine whether simple loss of positive charge at surface exposed Lys residues may promote SOD1 aggregation, we introduced mutations at four positions encoding Lys positions (K30V, K36T, K75A, K91G). The mutations introduced amino substitutions found at other positions in ALS (e.g. K30V similar to A4V, K36T similar to I113T, K75A similar to G93A, and K91G similar to D101G). As described in the Introduction, we used a visual assay to assess aggregation in which the mutations were introduced into fusion proteins of SOD1 and YFP. The use of the visual assay helps to control for variations in expression levels as fluorescence intensity can be used as a gauge for relative expression. As described in Methods, three transient transfections were performed for each construct and representative images were captured at 24 and 48 hours. For quantitative analysis of aggregate formation, cells expressing the fusions proteins were identified by fluorescence and scored for the presence of inclusion-like structures. Similar to our previous work with this approach [61,62], a subset of cells adopted a condensed, rounded, morphology and these cells were not scored because we could not easily discern whether they contained inclusions. These condensed cells do not display markers associated with cell death [TUNEL labeling or dimeric Ethidium bromide uptake [61]], and the basis for their appearance is unknown.

**FIGURE 1.**
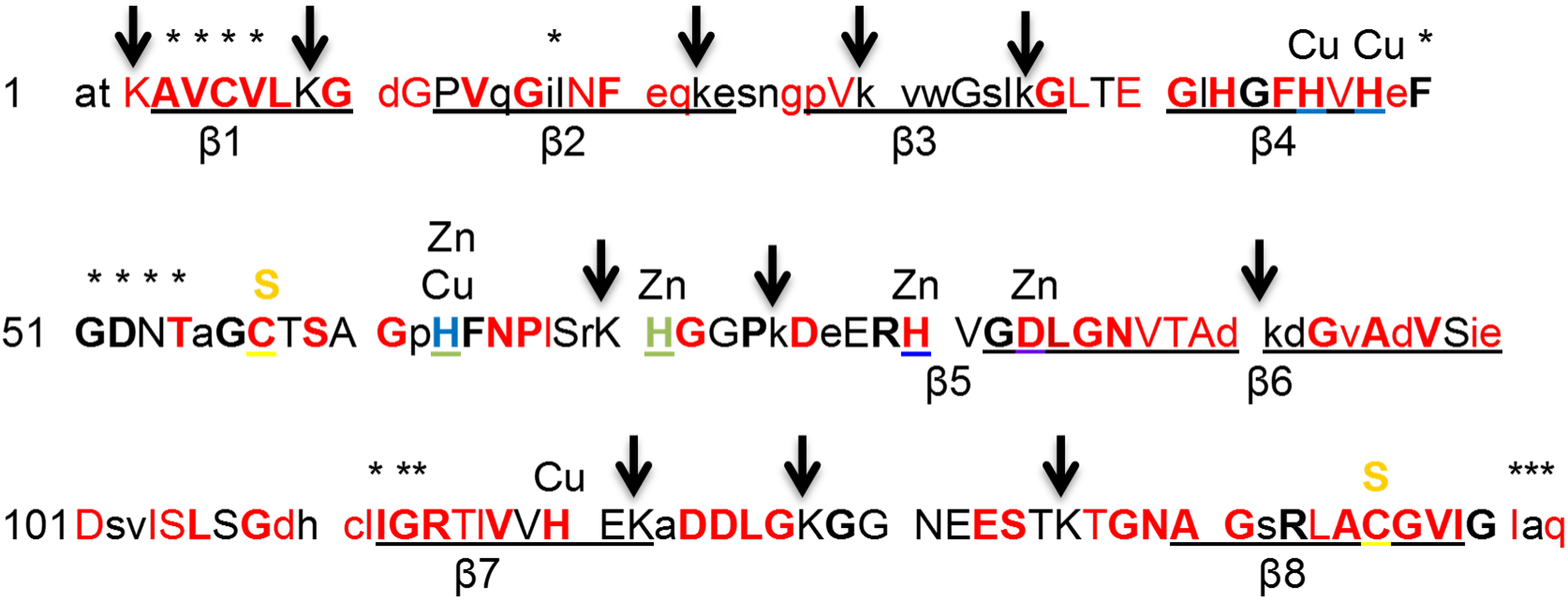
Location of mutations in SOD1 that have been identified in patients with ALS {http://alsod.iop.kcl.ac.uk}. Red letter font indicates a position where a point mutation has been identified in ALS patients. Bold indicates highly conserved amino acids across multiple species. Capitalized indicates conserved in mammals. Asterisks mark amino acids near the dimer interface. Arrows mark the position of Lys residues. Charged residues with mutations associated with ALS {http://alsod.iop.kcl.ac.uk}: D11Y; D76Y,V; D83G; D90A,V; D96V,N; D101G,N,Y,H; D109Y; D124V,G; D125H; E21K,G; E40G; E49K; E100G,K; E133d,V; H43R; H46R,D; H48Q; H80R; H120L; R115G,C; K3E. Positions lacking mutations at time of writing {http://alsod.iop.kcl.ac.uk}: D52, D92, E24, E77, E121^‡^, E132^†^, H63, H71, H110, R69, R79, R143, K9, K23, K30, K36, K70, K75, K91, K122^‡^, K128^‡^, K136^‡^; ^†^insertion frame-shift mutation at this site; ^‡^can be lost by early termination mutations.

Focusing on cells with a scorable morphology, mutations at four different Lys positions (K30V, K36T, K75A, K91G) produced very few cells with inclusions at either time point (Fig. 2; Table 1). Immunoblotting of cell lysates confirmed that the expression levels of these fusion proteins was similar to that of the positive control G93A-SOD1:YFP variant (Fig. 3). In addition to assessing the effect of mutations on SOD1 aggregation, we examined whether the mutations may produce more subtle effects on folding, using the binding of C4F6 antibody as an assay [45,61,68,69]. The epitope for C4F6 has been partially defined as including amino acids D90, D92, G93,and D96 of the protein segment 90DKDGVAD96 [45,69]. When C4F6 is used in immunocytochemistry of fixed cells transiently over-expressing mutant SOD1, cells expressing many different fALS mutants show much stronger reactivity than cells expressing WT protein [45,61]. Importantly mutations at residues located relatively far from the C4F6 binding site, such as the A4V mutation, enable the binding of C4F6 [61]. Thus, C4F6 can be used as a sensitive detector of changes in SOD1 conformation when used in immunocytochemistry. As compared to cells expressing G93A-SOD1, cells expressing K30V, K36T, K75A, or K91G variants showed a lower frequency of C4F6 immunoreactivity (Fig. 4), but not as low as cells expressing WT SOD1:YFP (Table 1). The K91G mutant produced the lowest frequency of C4F6 binding, but because this residue is within the epitope recognized by C4F6 (second amino acid) we were uncertain as to whether the mutation lowered antibody affinity.

**FIGURE 2.**
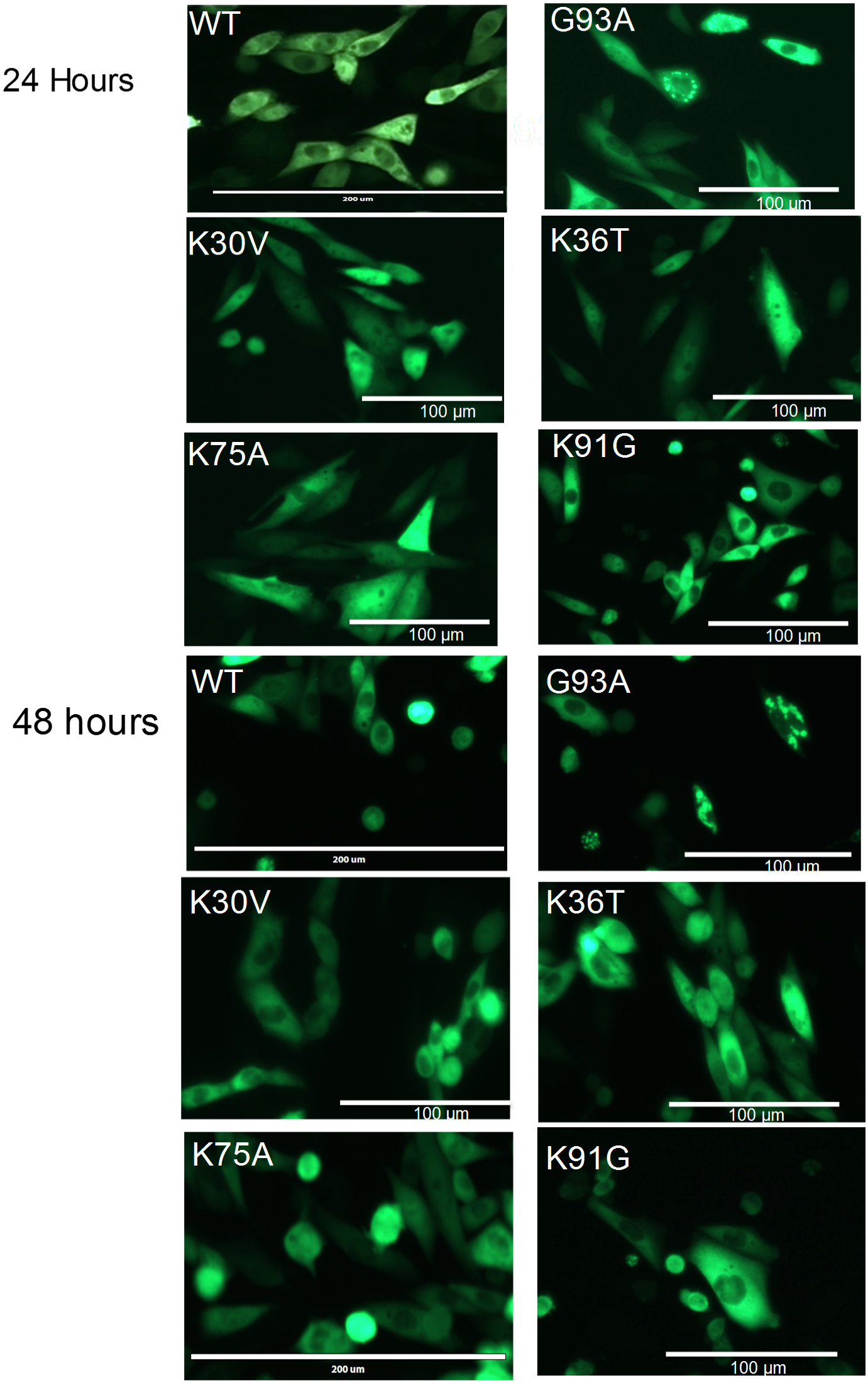
Cell assay of aggregation by SOD1 variants encoding mutations at lysine residues. CHO cells were transfected with plasmids for each SOD1:YFP variant and representative pictures were taken at 24 and 48 hours post-transfection. G93A-SOD1:YFP provides a positive control for inclusion formation and WT provides a negative control for inclusion formation. The percentage of cells expressing these mutants (K30V, K36T, K75A, or K91G) that developed inclusions was far lower than the positive control G93A variant (Table 1).

**FIGURE 3.**
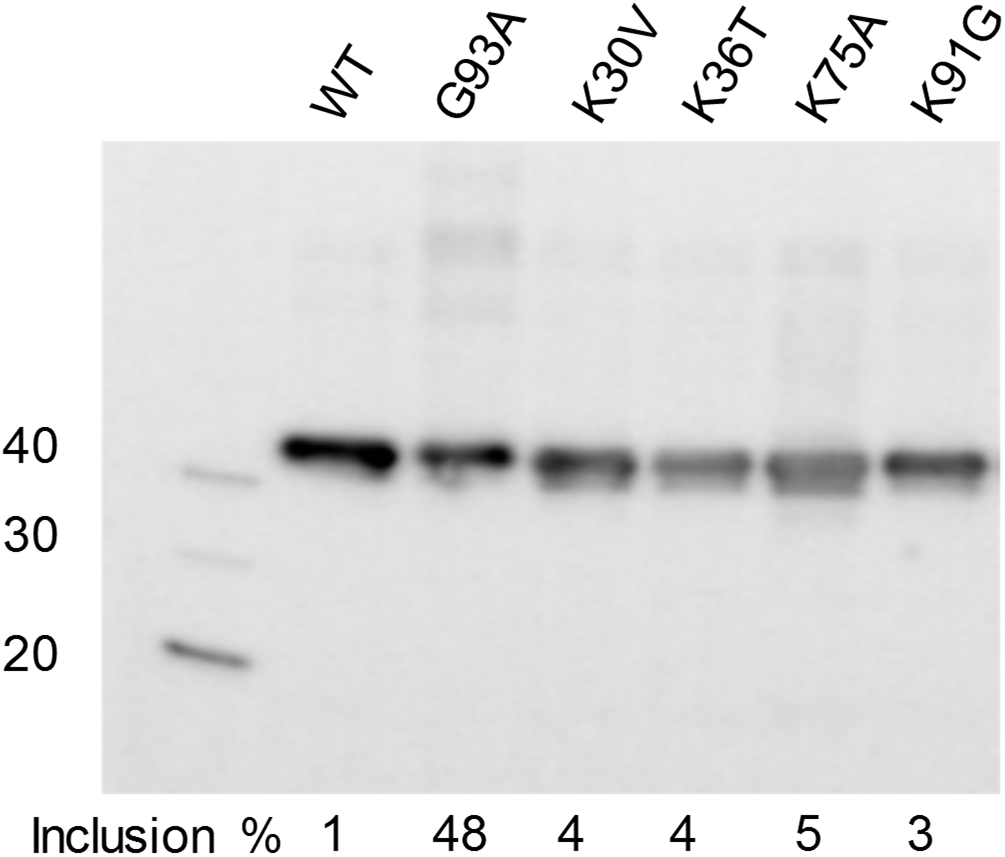
Immunoblot analysis of cells expressing WT and mutant SOD1 fused to YFP. CHO cells were transfected with plasmids for each SOD1:YFP variant and harvested at 24 hours for immunoblot analysis with antibody to SOD1 (see Methods). Despite similar levels of expression, the percentage of cells expressing these mutants (K30V, K36T, K75A, or K91G) that developed inclusions was far lower than the positive control G93A variant (see Table 1). The image shown is representative of 3 independent immunoblotting experiments. At the level of expression and exposure shown here, endogenous CHO SOD1 is not visible.

**FIGURE 4.**
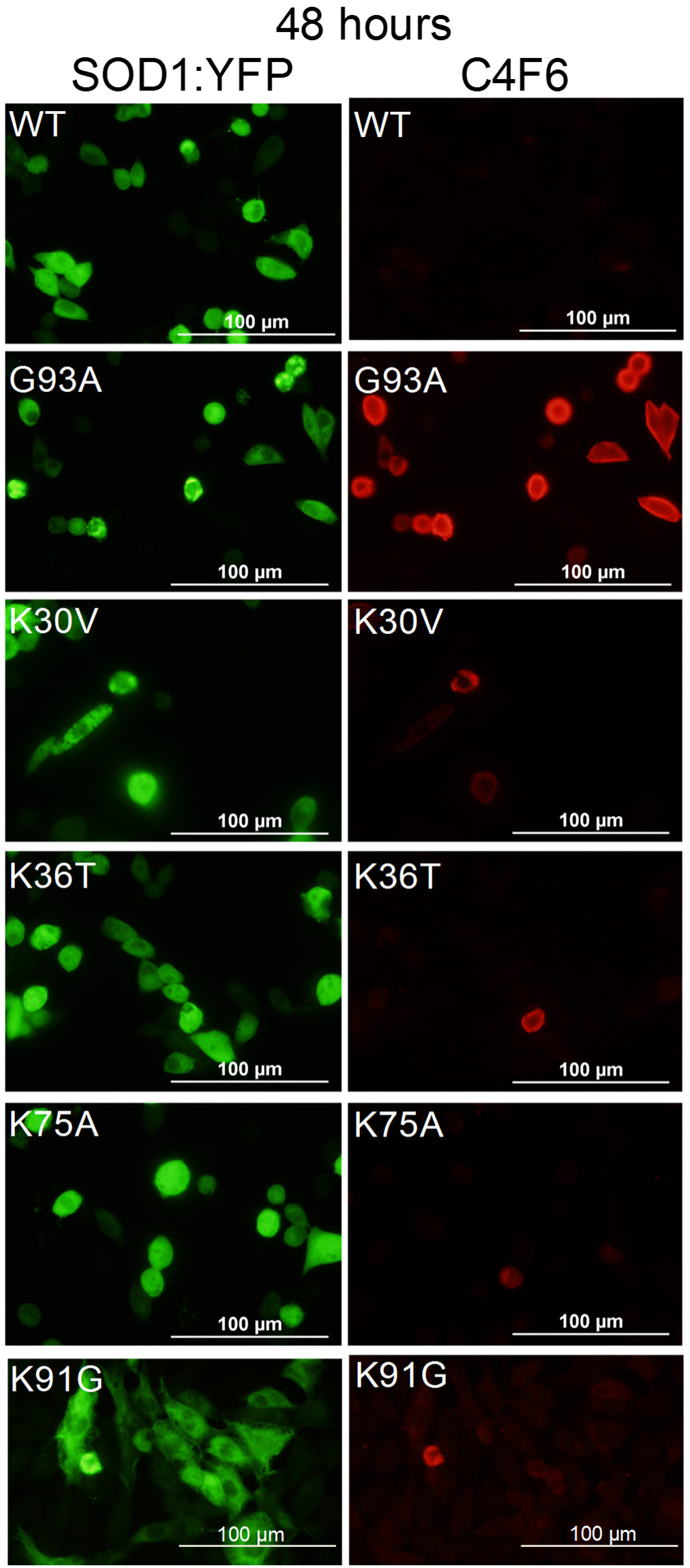
Analysis of C4F6 binding by SOD1 variants encoding mutations at Lys residues. CHO cells were transfected with plasmids for each SOD1:YFP variant and after 48 hours the cells were fixed and immunostained as described in Methods. Cells transfected with WT-SOD1:YFP serve as a negative control and cells transfected with G93A-SOD1:YFP serve as a positive control. Each transfection was repeated at least 3 times and at least 3 fields of view were counted for cells expressing the YFP fusion protein that were also labeled with C4F6 (Table 1).

**Table 1.** Quantification of inclusion frequency and C4F6 reactivity.

| SOD1 Variant | # cells counted (24 hr) | # cells with inclusions (24 hr) | % cells with inclusions (24 hr) | # cells counted (48 hr) | # cells with inclusions (48 hr) | % cells with inclusions (48 hr) |
| --- | --- | --- | --- | --- | --- | --- |
| WT | 492 | 3 | $\leq 1\%$ | 742 | 6 | 1% |
| G93A | 380 | 196 | 52% | 503 | 243 | 48% |
| K30V | 431 | 15 | 3.4% | 618 | 25 | 4% |
| K36T | 550 | 13 | 2.4% | 736 | 29 | 4% |
| K75A | 377 | 9 | 2.4% | 530 | 26 | 5% |
| K91G | 207 | 1 | $\leq 1\%$ | 123 | 4 | 3% |
| SOD1 Variant | # cells counted (24 hr) | # cells with C4F6 reactivity (24 hr) | % cells with C4F6 reactivity (24 hr) | # cells counted (48 hr) | # cells with C4F6 reactivity (48 hr) | % cells with C4F6 reactivity (48 hr) |
| WT | 114 | 13 | 11% | 415 | 9 | 2% |
| G93A | 419 | 370 | 88% | 340 | 307 | 90% |
| K30V | n.d. | n.d. | n.d. | 145 | 31 | 21% |
| K36T | n.d. | n.d. | n.d. | 164 | 14 | 9% |
| K75A | n.d. | n.d. | n.d. | 96 | 15 | 16% |
| K91G | n.d. | n.d. | n.d. | 171 | 10 | 5% |

To define the C4F6 epitope further, we examined its reactivity to a panel of SOD variants by immunoblotting. The C4F6 antibody was raised against recombinant human G93A SOD1 [68]. In immunoblot analysis of a panel of ALS variants at the G93A position (G93A, G93C, G93D, G93R, G93V), it was clear that the antibody exhibited the greatest avidity for G93A, with moderate levels of reactivity to WT (G93) protein (Fig. 5). To determine whether K91 was critical in C4F6 binding, we generated a panel of mutations at Lys 91 (K91G, K91N, K91R, and K91E). The K91G variant was poorly reactive (Fig. 5). However, through the introduction of multiple mutations at K91, we determined that multiple mutations at K91 could be introduced without disrupting C4F6 binding (Fig. 5). To assess the importance of K91 in C4F6 binding in fixed cells, we generated a double mutation of K91E/G93A fused to YFP and expressed it in CHO cells. At 48 hours post-transfection, 95% of cells expressing the K91E/G93A variant showed C4F6 reactivity (Fig. 6). Collectively, these data indicate that the Lys at position 91 is not required for C4F6 binding. Thus, the lack of binding of the K91G variant by C4F6 likely reflects a consequence of a change in the flexibility of this segment of SOD1 that lowers C4F6 affinity. Overall, our data show that mutations at surface exposed Lys residues K30, K36, K39, K75, and K91 do not produce sufficient conformational changes to evoke aggregation, but most of these mutations evoke moderate C4F6 binding (Table 2).

**FIGURE 5.**
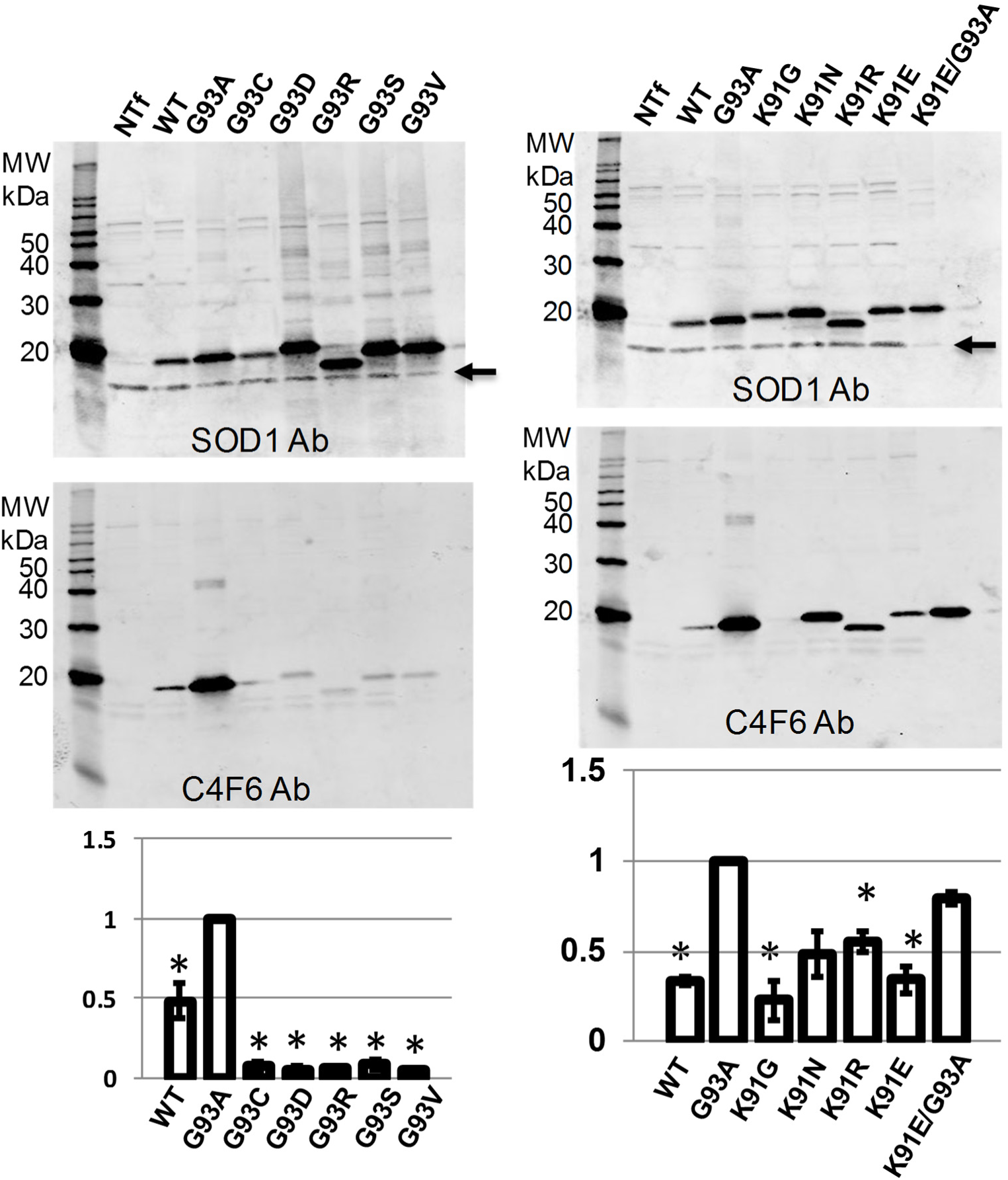
Immunoblot of cell lysates from CHO cells transfected with SOD1 variants encoding fALS mutations at Gly 93. The cell expressing these SOD1 variants (untagged) were harvested at 48 hours post-transfection, lysed, and analyzed by immunoblotting with the C4F6 and SOD1 antibodies as described in Methods. The binding of C4F6 to denatured SOD1 was greatly reduced if G93 was mutated to other residues that cause fALS, including G93C, G93D, G93R, G93S, and G93V. When position 93 is Gly, the affinity of C4F6 is inherently weak but apparently can be augmented by conformational changes that modulate the accessibility of the epitope. The data were quantified from 3 independent transfection experiments as described in Methods. All mutants tested were less reactive to C4F6 than the G93A variant (^*^ p<0.05 G93A versus all other variants, T-Test, 2 tailed, unequal variance). The G93C, G93D, G93R, G93S, and G93V variants were all less reactive than WT SOD1 to C4F6 (^**^ p<0.05, T-Test, 2 tailed, equal variance). The arrow marks the positon of endogenous CHO SOD1.

**FIGURE 6.**
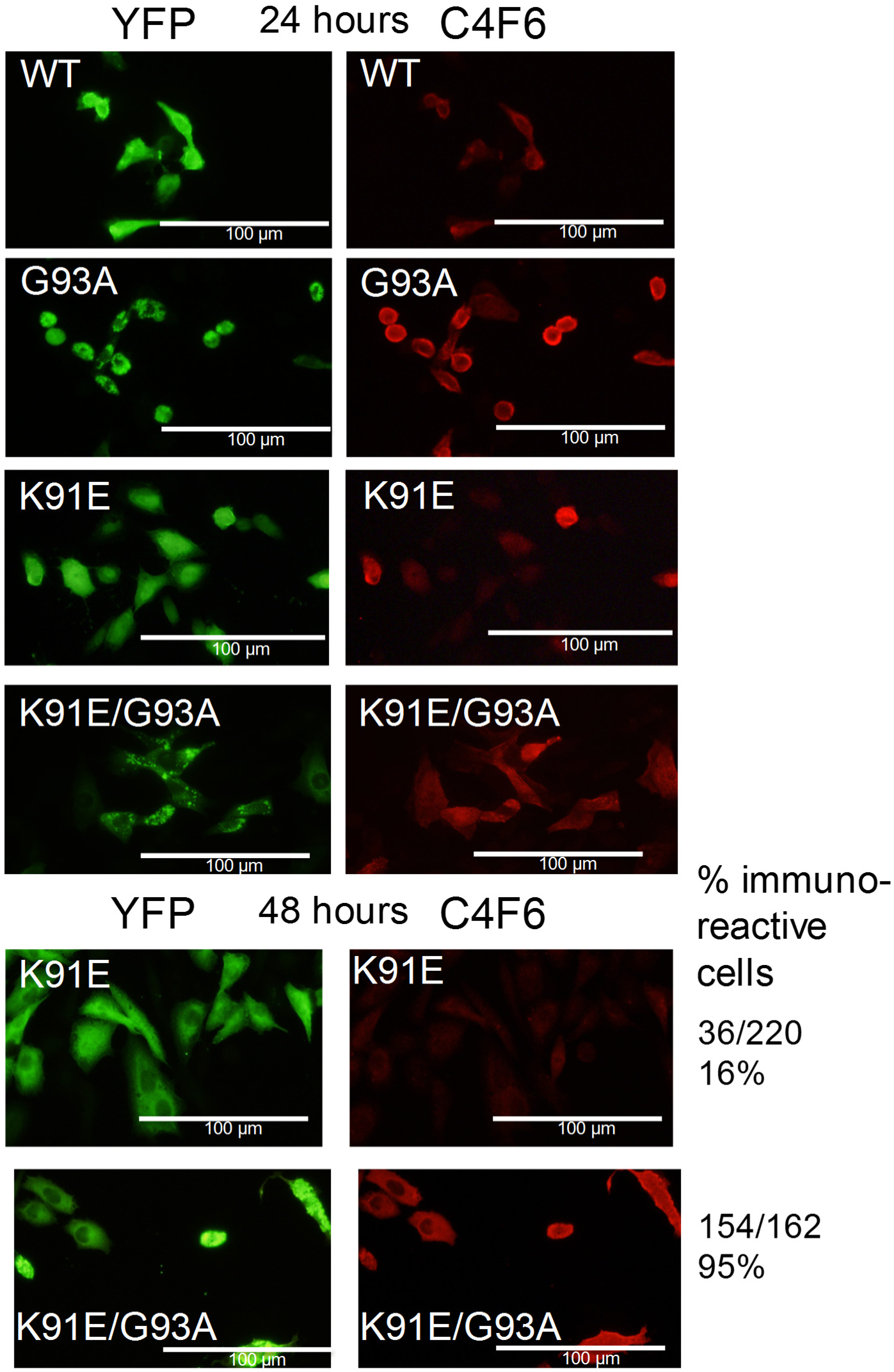
Analysis of C4F6 binding by SOD1 variants encoding an experimental K91E mutation in G93A hSOD1. CHO cells were transfected with plasmids for each SOD1:YFP variant and after 24 or 48 hours the cells were fixed and immunostained as described in Methods. Cells transfected with WT-SOD1:YFP serve as a negative control and cells transfected with G93A-SOD1:YFP serve as a positive control (see Fig. S4 for an example of 48h for each). Each transfection was repeated at least 3 times and at least 3 fields of view were counted for cells expressing the YFP fusion protein that were also labeled with C4F6 (Table 2). Mutation of Lys 91 to Glu does not markedly diminish the binding of C4F6 antibody.

## DISCUSSION

Using a visual read-out for mutant SOD1 aggregation and an antibody based approach to detect misfolding, we examined relationships between loss-of-charge at positively charged positions in hSOD1 and the propensity to misfold and aggregate. Mutations in lysine residues of hSOD1 are rare in fALS patients, and we generally found that substitution mutations in lysine were well tolerated by the protein in terms of propensity to aggregate. However, mutations at two lysine residues (K30, K75) modestly increased the frequency of cell reactivity to C4F6 (Table 2), suggesting that these mutations disturbed protein conformation to some degree. Based on the behavior of the mutant SOD1:YFP fusion proteins studied here, we conclude that a simple loss of positive charge at solvent exposed Lys residues is not necessarily sufficient to alter the inherent propensity of SOD1 to aggregate.

Our method for assessing aggregation involved visual detection of cytoplasmic inclusion formation in mammalian cells using a platform of fusing variants of SOD1 to YFP. In prior studies we have demonstrated that cells induced to express mutant SOD1:YFP fusion genes by transient transfection produce fluorescent inclusions that is accompanied by an increase in the level of detergent insoluble SOD1:YFP [61]. Wild-type or mutant SOD1:YFP fusions proteins that show a diffuse distribution in the cytosol are readily released into cell medium by treatment with saponin, whereas the inclusion structures (when present) remain with the cell after saponin [61]. Collectively, these prior studies indicate that a diffuse distribution of SOD1:YFP in cell cytosol is associated with high mobility and solubility. Thus, the lack of inclusion formation by these Lys mutants of SOD1 fused to YFP suggests strongly that they can achieve a soluble, relatively native, conformation.

We complemented the visual screen for aggregation with an immunological screen in which we determined whether mutations in SOD1 increased reactivity to the monoclonal antibody C4F6. This antibody was raised against purified recombinant G93A protein [68]. We have previously demonstrated that multiple fALS mutant SOD1:YFP fusion proteins are reactive with the C4F6 antibody [61], which has been described as being specific for misfolded mutant SOD1 [45,68]. In cells expressing G93A SOD1, ~90% of the cells are C4F6 reactive. Notably, the C4F6 antibody demonstrates higher avidity for G93A variant SOD1 over WT, or any other FALS substitution at Gly 93, in immunoblots of SDS-denaturing PAGE. This finding implies that the Ala at position 93 is an integral component of the epitope and thus, the high degree of C4F6 immunoreactivity in fixed cells expressing the G93A variant may be due to both loss of native structure and higher avidity for SOD1 when position 93 is Ala. Still, as compared to WT-SOD1:YFP, which rarely produces cells that are C4F6 reactive (Table 1), some of the mutants tested (K30V and K75A) produced moderate to modest frequencies of C4F6 reactive cells (Table 1). These data suggest that the mutations at surface exposed Lys residues that we have analyzed may modestly alter folding without inducing aggregation.

### Conclusions

Although the nature of the toxic form of mutant SOD1 in fALS remains imprecisely defined, it is clear that one consequence of disease-causing mutations in hSOD1 is to destabilize normal structure in a manner that facilitates aberrant homotypic self-assembly into higher order structures [50,70]. In the present study, we asked whether mutations that reduce positive charge would potentially induce aggregation by reducing repulsive forces as SOD1 aberrantly assembles into aggregates. Our data indicate that mutations at Lys 30, 36, 75, and 91 that would reduce surface positive charge are relatively well tolerated. Our data may explain why mutations at these conserved Lys residues are rarely found in ALS patients.

## EXPERIMENTAL PROCEDURES

### Generation of mutant SOD plasmids

-Human SOD1 cDNA’s with mutations described in this study were generated using oligonucleotide primers encoding the desired mutation with Quick-Change mutagenesis kits. The single mutants were made using pEF.BOS vectors [71] that encode WT human SOD1 (WT-hSOD1) as the template [51]. The protocol used to make mutations used a modified PCR strategy with primers encoding specific mutations and pEF-BOS-WT SOD1 or pEF-BOS-WT SOD1:YFP plasmids as the template. The PCR reaction used Platinum *Pfx* polymerase (Invitrogen/ThermoFisher, Waltham, MA) and 2X Pfx buffer concentration to accommodate the large plasmids that were amplified. The PCR reaction products were digested with Dpn1 to remove template and then transformed into NEB-10β competent cells (New England Biolabs, Ipswich, MA) following standard protocols. Large scale preparations of plasmid DNA for transfection were prepared by CsCl gradient purification. The SOD1 and SOD1:YFP coding sequence of all plasmids was verified by DNA sequence analysis.

### Transient Transfections

-Plasmid DNA encoding the mutant SOD1:YFP cDNAs were transiently transfected into Chinese hamster ovary (CHO) cells. The day before the transfection, the CHO cells were split into 60-mm poly-D-lysine-coated dishes (1 plate for each DNA construct). Upon reaching 95% confluency, cells were transfected with Lipofectamine^™^ 2000 (Invitrogen/ThermoFisher). The cells were then incubated at 37°C in a CO2 incubator for 24 hours at which time images of random fields of view at 20x and 40 x magnification were captured using an AMG EVOS_fl_ digital inverted microscope for fluorescence. The cells were returned to the incubator for 24 hours before images were captured again.
The transient transfections were repeated at least 3 times for each construct. The images from multiple transfections were analyzed and the following objects were counted in a blinded fashion, cells showing YFP fluorescence, fluorescent inclusions, and fluorescent cells with condensed morphology.

### Immunocytochemistry

The day before transfection, CHO cells were plated onto poly-D-lysine coated glass coverslips. 24 or 48 hours following transient transfection with pEF-BOS-SOD1 constructs, cells were fixed with 4% paraformaldehyde for 10 minutes at room temperature. Following washing with 1X PBS, cells were perforated with ice-cold methanol. Non-specific binding was blocked with 5% normal goat serum for 1 hour at room temperature. Human SOD1 was identified using the human-specific hSOD1 rabbit antibody [72] and misfolded SOD1 with the mouse monoclonal antibody C4F6 [68] diluted 1:1000 in 3% normal goat serum and incubated at 4 degrees Celsius overnight. Following PBS washes, cells were incubated with goat anti-rabbit AlexaFlour-488 and goat anti-mouse AlexaFlour-568 (Invitrogen) at 1:2000 in 3% normal goat serum with 1:2000 dilution of DAPI at room temperature for 1 hour. Cells were washed with PBS and mounted to glass slides (Fisher) with Aqua-Poly/Mount coverslipping medium (Polysciences, Inc, Warrington, PA). Fluorescent microscopy images were captured on an Olympus BX60 epifluorescence microscope. Most images were captured at 20x.

### Immunoblotting

- At 48 hours post-transfection, CHO cells were washed from the plate in 1X PBS, and then centrifuged at 3000xg rpm for 5 minutes before resuspension in 1X PBS with protease inhibitor cocktail (Sigma, St. Louis, MO, USA). Cells were disrupted with a probe sonicator for 10 seconds, and protein concentrations determined with BCA assay (Pierce/ThermoFisher, Waltham, MA). 5μg of total protein was loaded and separated through an 18% TG-SDS PAGE gel (Invitrogen/ThermoFisher) and transferred to nitrocellulose membrane. The membranes were incubated in Odyssey blocking solution (Li-Cor, Lincoln, Nebraska) as directed by the manufacturer and then primary antibodies (SOD1 whole protein rabbit polyclonal antibody [73] and C4F6 monoclonal antibody) were incubated at 1:2000 overnight in Odyssey blocking solution. The membranes were washed in 1X TBS-T, and probed with goat anti-rabbit IRDye-680RD and goat anti-mouse IRDye-800CW near-infrared-labeled secondary antibodies (Li-Cor). Images were captured using an Odyssey imaging system (Li-Cor) and densitometry analysis was performed using ImageJ (NIH, Bethesda, MD). The values for the intensity of SOD1 bands were normalized to G93A SOD1 protein and the data were graphed using Excel.

## ACKNOWLEDGEMENTS

– We are grateful for helpful advice from Drs. P. John Hart, Julian Whitelegge, David Eisenberg, and Joan S. Valentine. We thank undergraduate students Aron Workman, Adam DeBossier, and Kinaree Patel for their contributions to this study. The authors declare they have no competing interests.

## FUNDING

This work was supported by a grant from the National Institutes of Neurological Disease and Stroke (P01 NS049134) and by the University of Florida.

^**^These authors made equal contributions to this manuscript.

The abbreviations used are: SOD1, superoxide dismutase 1; ALS, amyotrophic lateral sclerosis; fALS familial amyotrophic lateral sclerosis; YFP, yellow fluorescent protein

